# Personality homophily affects male social bonding in wild Assamese macaques (*Macaca assamensis*)

**DOI:** 10.1101/520064

**Authors:** Anja Ebenau, Christoph von Borell, Lars Penke, Julia Ostner, Oliver Schülke

## Abstract

Animal social bonds are defined as stable, equitable and strong affiliative and cooperative relationships similar to human friendships. Just as human friendships, social bonds are thought to function as alliances that generate adaptive benefits via support in critical situations. In humans, similarity in many sociodemographic, behavioural and intrapersonal characteristics leads to trust and is predictive of friendships. Specifically, personality homophily, that is the tendency of individuals to form social bonds with others who have a similar personality, may increase predictability and facilitate trust and reciprocity among partners with compatible behavioural tendencies. While evidence for social bonding in nonhumans is accumulating, far less is known about its predictors. Here, personality homophily effects on the formation and maintenance of social bonds are shown in twenty-four wild male Assamese macaques (*Macaca assamensis*), at Phu Khieo Wildlife Sanctuary, Thailand. Dyadic bond strength increased with increased similarity in the trait Connectedness (i.e. frequent and diverse neighbours in 5m proximity and pronounced social tolerance, as high rates of friendly approaches to and by others). To differentiate whether homophily indeed predicted bond formation or whether bonded males’ personalities became more similar over time, we tested the stability of the connectedness traits in a subset of immigrating males that had to form new bonds. Connectedness in these males remained stable suggesting that males do not adapt their personality to their partner. Our results support the idea of a shared evolutionary origin of homophily as a partner choice strategy in human and non-human animals. The main selective advantage of personality similarity in animal social bonds may result from a more reliable cooperation among individuals with similar cooperative behavioural tendencies.

## INTRODUCTION

In mammals and birds, social bonds are defined as stable, equitable and strong affiliative relationships similar to human friendships, and like friendships are thought to function as alliances that generate adaptive benefits via support in critical situations (Brown & Brown, 2006; Cheney, 2011; Curry & Dunbar, 2013; DeScioli & Kurzban, 2009; Ostner & Schülke, 2014, 2018; Schino, 2007; Silk, 2007). Bond strength promotes coalition formation (e.g., Berghänel, Ostner, Schröder, & Schülke, 2011; Connor, Heithaus, & Barre, 2001; Gilby et al., 2013; Perry, Barrett, & Manson, 2004; Watts, 2002; Young, Majolo, Schülke, & Ostner, 2014) and enhances cooperative success, possibly through increased trust in a bonded ally (across a wide range of taxa in birds and mammals: Braun & Bugnyar, 2012; Engelmann & Herrmann, 2016; Majolo et al., 2006; Marshall-Pescini, Schwarz, Kostelnik, Virányi, & Range, 2017; Massen, Ritter, & Bugnyar, 2015; Molesti & Majolo, 2016; Olson & Spelke, 2008; Wood, Kim, & Li, 2016). In risky situations, when an individual has to choose with whom to cooperate, social bonds spare situational judgement and cognitive effort of assessing partner quality and honesty of signals, since they reduce uncertainty about the partner’s response (Cronin, 2012; Molesti & Majolo, 2016; Noë, 2006; Schino & Aureli, 2009). According to standard evolutionary models, partner choice mechanisms are key to initiate and maintain cooperative behaviours, and can lead to the formation of differentiated social relationships from weak ties to social bonds in animal groups (Campennì & Schino, 2014; Noë, 2006; Schino & Aureli, 2016).

Partner choice for the formation of social bonds may be guided by homophily, that is the tendency of individuals to form ties with similar others (McPherson, Smith-Lovin, & Cook, 2001), as it may enhance predictability and trust in potential bond partner (Dunbar, 2018; Massen & Koski, 2014; Weinstein & Capitanio, 2012). Animal and human social structure in terms of spatial or socio-ecological associations partly results from assortment by age, sex, kinship, body size, reproductive state, or genotype (e.g., Fowler, Settle, & Christakis, 2011; Fu, Nowak, Christakis, & Fowler, 2012; McPherson et al., 2001).

Fitness-related advantages of choosing self-similar bond partners may arise from a shared mode of communication and more efficient coordination and cooperation (Fu et al., 2012; Noë, 2006). In theoretical models, homophily enhances the evolution of cooperation and facilitates the spread of cooperation in human and nonhuman animal networks (Antal, Ohtsuki, Wakeley, Taylor, & Nowak, 2009; Chiang & Takahashi, 2011; Nowak, Tarnita & Antal, 2010; Riolo, Cohen, & Axelrod, 2001; Rivera, Soderstrom, & Uzzi, 2010; Voelkl & Kasper, 2009).

In humans, similarity in many sociodemographic, behavioural and intrapersonal characteristics, as well as sharing values, leads to trust and predicts friendships more than dissimilar characteristics (Bahns, Crandall, Gillath, & Preacher, 2016; Curry & Dunbar, 2013; Kandel, 1978; McPherson et al., 2001; Selfhout, Branje, & Meeus, 2007; Ziegler & Golbeck, 2007). Trust also plays a crucial role in forming and maintaining relationships in nonhuman animals, particularly in non-kin (Dunbar, 2018; Engelmann & Herrmann, 2016; Massen & Koski, 2014; Massen et al., 2015). Chimpanzees selectively trust bonded partners (Engelmann & Herrmann, 2016), suggesting that trust in reciprocity is not unique to humans, but has deeper evolutionary roots (Engelmann, Herrmann, & Tomasello, 2015). In male Barbary macaques the probability that a bystander of an aggressive conflict rejects a recruitment for help decreased with the strength of the affiliative relationship between the bystander and the recruiter (Young et al., 2014), i.e. the individual in need can trust that bonded partners will provide support.

Trust and reciprocity may be facilitated specifically via homophily in personality (Hampson, 2011; Massen, 2017; Massen & Koski, 2014; Scarr & McCartney, 1983). Personality is defined as inter-individual differences in behaviour, affect and cognition that are relatively consistent across time and (Réale, Reader, Sol, McDougall, & Dingemanse, 2007). Personality homophily has been found in human spouses (e.g., Byrne, 1997; Klohnen & Luo, 2003; Youyou, Stillwell, Schwartz, & Kosinski, 2017) and improves reproductive success in monogamous rodents, birds, and fish (Ariyomo & Watt, 2013; Dingemanse, Both, Drent, & Tinbergen, 2004; Gabriel & Black, 2012; Rangassamy, Dalmas, Féron, Gouat, & Rödel, 2015; Schuett, Dall, & Royle, 2011). Similarity in certain personality traits is associated with the strength of social bonds in chimpanzees (Massen & Koski, 2014), higher-quality relationships in capuchin monkeys (Morton, Weiss, Buchanan-Smith, & Lee, 2015), relationship stability from one year to the next in juvenile rhesus macaques (Weinstein & Capitanio, 2012) and pairing-success of adult rhesus macaques in a laboratory setting (Capitanio, Blozis, Snarr, Steward, & McCowan, 2015). Beyond dyadic relationships, group-level similarity in personality traits facilitates cooperation among all group members in cooperative-breeding common marmosets (Koski & Burkart, 2015).

Friends with similar personalities may perceive, interpret, and react to the world around them in a similar way (neuronal homophily; Parkinson, Kleinbaum, & Wheatley, 2018). Friends share dispositions and agree on values, opinions and activities, which may trigger a positive affective response that increases enjoyment of each other’s company, and strengthens the self-concept (Baumeister & Leary, 1995; Campbell, Sedikides, Reeder, & Elliot, 2000; Clore & Byrne, 1974; Hampson, 2011; Nelson, Thorne, & Shapiro, 2011; Nelson et al., 2011; Selfhout et al., 2010). Personality similarity among friends may further reduce uncertainty during acquaintanceship and enhances predictability by increasing the ease and clarity of communication (Berger & Calabrese, 1975; Neyer, Banse, & Asendorpf, 1999; Selfhout et al., 2010; van Zalk & Denissen, 2015). With respect to the “Big Five” personality model (Digman, 1990; John, Srivastava, & Pervin, 1999), friends are mostly found to be similar in two dimensions (e.g., Blaz, 1983; Caspi, Roberts, & Shiner, 2005; Feiler & Kleinbaum, 2015; Jensen-Campbell et al., 2002: extraversion, a dimension capturing variation in activity, sociability, positive emotionality, risk seeking and assertiveness, and agreeableness which describes variation in being kind and considerate, empathic, prosocial and cooperative (van Aken & Asendorpf, 2018). Given the potentially shared evolutionary history of social bonds and human friendships (Baumeister & Leary, 1995; Seyfarth & Cheney, 2012; Silk, 2002), and the fact that shared neural and physiological mechanisms underlie social behaviours in humans and other animals (Brent, Chang, Gariépy, & Platt, 2014; Chang et al., 2013; Dunbar, 2010; Meunier, 2018), it has been proposed that homophily in human social partner choice has a biological basis (Apicella, Marlowe, Fowler, & Christakis, 2012; Bahns et al., 2016; Fu et al., 2012; Massen & Koski, 2014; Parkinson et al., 2018).

Here we investigated whether patterns of affiliation correspond to homophily in personality traits in wild male Assamese macaques. Apart from an unpublished PhD thesis (Tkaczynski, 2017) these studies all used captive animals and assessed personality either with behavioural or with trait rating (i.e. questionnaire) data. We add ecological validity by studying wild animals. Male Assamese macaques are particularly well-suited for this study, because males change groups several times during their life (Ostner, Vigilant, Bhagavatula, Franz, & Schülke, 2013), and because males in the study population form differentiated social bonds that convey fitness benefits via increased paternity success (Kalbitz, Ostner, & Schülke, 2016; Schülke, Bhagavatula, Vigilant, & Ostner, 2010).

Instead of predicting homophily for a particular personality dimension, we followed an explorative approach and expected to find homophily in any of the five personality traits we defined for these males, namely Connectedness, Aggressiveness, Sociability, Vigilance, and Confidence (Ebenau, Penke, Ostner, & Schülke, under review). In humans the social personality traits extraversion and agreeableness are similar among friends, but other traits may affect social partner choice as well: bonded partners are more similar in boldness in chimpanzees (Massen & Koski, 2014) and traits like aggressiveness may be more relevant in some species as it is shaping the social style in macaques (Adams et al., 2015). As closely bonded individuals pull each other to similar ranks via support in agonistic interactions with the benefits of increased access to food and mates (Chapais, 1995; Schülke et al., 2010), we expected and therefore controlled for an effect of dominance rank difference on dyadic social bond measures. We expect that similarity in personality predicts bond formation. To rule out that this correlation results from bonded partners adapting their personalities over time, we assess personality stability in males changing social groups during the study period, which is accompanied by changing affiliation partners.

## METHODS

Fieldwork was conducted in the Phu Khieo Wildlife Sanctuary (PKWS: 16°5’–35’N, 101°20’–55’E) which is part of the ca. 6500 km^2^ interconnected and well-protected Western Isaan forest complex in north-eastern Thailand (Borries, Larney, Kreetiyutanont, & Koenig, 2002). The study area is covered by hill evergreen forest and harbours a diverse community of large mammals and predators (Borries et al., 2002) indicative of very low levels of human disturbance. The field site was established in 2005, study subjects lived in four fully habituated groups, and were followed from April 2014 (ASM and AOM group) or October 2014 (ASS and AOS group) through March 2016. Group sizes at the beginning of behavioural data collection are shown in Table A1.

### Personality assessment

We applied a multi-method approach based on analyses of trait ratings (TR) and behavioural codings (BC), which allowed for testing construct validity of the quantified personality structures (for details see Ebenau et al., under review). In brief, individuals were rated twice in 2015 and 2016 on the 54 item Hominoid Personality Questionnaire (HPQ; King & Figueredo, 1997; Weiss et al., 2009). Each adjective item was defined within the context of general behaviours common to primates. For example, ‘fearful’ was defined as “Subject reacts excessively to real or imagined threats by displaying behaviours such as screaming, grimacing, running away or other signs of anxiety or distress.” Data were processed by analysing rater performance, applying interrater-reliability (ICC; Shrout & Fleiss, 1979) with a cut-off criterion of > 0.4, and examining temporal stability from one year to the next. After data reduction, 43 adjective items were submitted to factor analysis, revealing four dimensions: Aggressiveness_TR_, Confidence_TR_, Activity_TR_ and Friendliness_TR_. To validate the rating data, behavioural codings were analysed for 24 adult males. Behavioural data were collected from April 2014 to March 2016 concurrently for behavioural personality assessment as well as for relationship measures, and is described in detail below. Eighteen temporally stable variables were reduced to four factors: Connectedness_BC_, Aggressiveness_BC_, Sociability_BC_ and Vigilance_BC_. Construct validity assessments suggested congruence between most dimensions from trait ratings and behavioural codings, with the exception of the Confidence_TR_ trait rating domain, which therefore was added as a fifth dimension to the behavioural coding personality constructs (for details see Ebenau et al., under review).

**Table 1.**
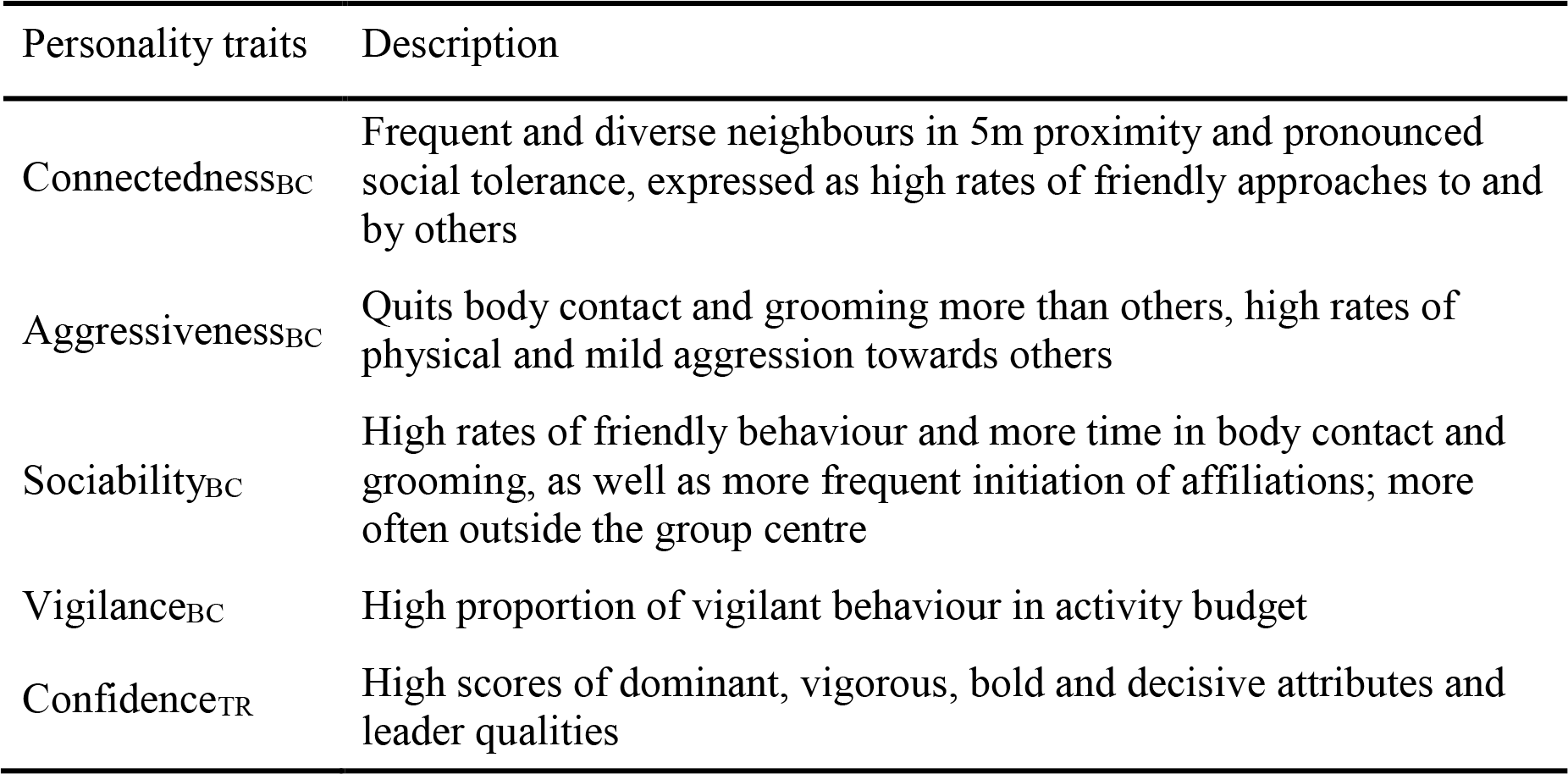
Summary of integrative personality constructs of Assamese macaques, derived from behavioural codingsbc and trait ratings_TR_.

### Behavioural data collection

We collected 4628 hours of focal animal observations (Altmann, 1974) from 24 adult males (mean per subject = 193 h; range = 86 h – 284 h) of the four study groups. These focal animals were included in the study, since they were present more than three months within one year of the two-year study period. Individuals were followed for 40 minutes with continuous recording of all approaches and departures within 1.5 m of the focal animal, and all affiliative and agonistic social interactions, with onset and termination for duration behaviours (e.g., approaches, body contact and grooming), as well as with directionality and the identities of interaction partners. Activity of the focal animal was recorded instantaneously at 2-minutes intervals. Every 10 min we recorded the identities of all individuals within a 5 m sphere around the focal animal. An effort was made to equally distribute observation time across individuals and time of the day. Quantitative behavioural data collected with a standardized ethogram were used to assess relationship strength.

### Dyadic relationship measure

For relationship assessment, we used data of two half-year periods with rather stable male group composition (October 2014–March 2015, October 2015–March 2016). Still, some adult males were absent for some time within these periods. We set two criteria and only included individuals, if they were either present in the group for at least half the time we spent with the group within the half-year period, or their observation hours did not fall below half the group mean within the half-year period. The remaining periods were too unstable to infer reliable relationship measures due to migration events as well as alpha male rank changes. Two of three adult males migrated from ASS into ASM group within the second year of observation, leaving only one adult male, thus, just one half-year period (October 2014–March 2015) was included for ASS group.

We used the dyadic sociality index (DSI; Silk, Cheney, & Seyfarth, 2013) to measure the strength of dyadic relationships, with frequencies and durations of correlated affiliative behaviours (mean τ_(rw,ave)_ = 0.491 ± 0.103), grooming, body contact and close proximity < 1.5 m. Since grooming frequencies between adult males tend to be quite low and to prevent inflation effects, we excluded grooming from the calculation when the average frequency across all dyads in a group was below 1.5. This was done for the second half-year period (October 2015–March 2016) for ASM and AOS group. For body contact and close proximity, we only included interactions longer than 10 seconds. Dyadic interaction rates and durations of overlaid behavioural states were subtracted from one another, and calculations were controlled for observation times of each partner. We calculated the index as follows:

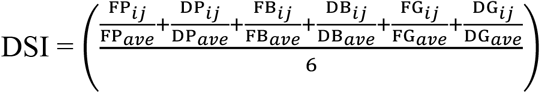

Here *ij* is the male-male dyad, *ave* is the group mean across all male-male dyads, **F** is the frequency and **D** the total duration of the behaviours: **P** as close proximity < 1.5 m, **B** as body contact and **G** as grooming. For a detailed description of dyadic CSI (i.e. DSI) calculation and its application in male Assamese macaques see Kalbitz et al. (2016). The index is a linear measure with a minimum of zero and a group mean of one, and increases with the strength of the affiliative relationship between two partners. Weak relationships are defined by values between zero and one, and values greater than 1 reflect stronger affiliative relationships (Silk, Alberts, & Altmann, 2006).

### Dominance rank

Male Assamese macaques can be ordered along a linear dominance hierarchy (Ostner, Heistermann, & Schülke, 2008), where higher-ranked individuals dominate all individuals of lower rank, thus all dyads have a dominant-subordinate relation. We calculated a dominance hierarchy from decided dyadic agonistic interactions as well as unprovoked submissive signals, e.g., silent-bared teeth (Ostner et al., 2008). Data on conflicts were recorded during continuous and ad libitum sampling for the same half-year period as the dyadic relationship measures. On average, we included in our analysis of dominance rank 13.7 and 16.3 interactions per individual in the two study periods respectively, which exceeds the value of 10 proposed for steep hierarchies (Sánchez-Tójar, Schroeder, & Farine, 2018). A winner/loser matrix of these interactions was used to calculate the standardized normalized David’s score (nDS) using DomiCalc (“compete” R-package; Schmid & de Vries, 2013). Due to group composition and alpha male rank changes we calculated an average rank for each period as a sum of hierarchical rank multiplied by the number of months the rank position was occupied divided by 6.

### Statistical analyses

We ran a linear mixed model (Baayen, 2008) to evaluate the effect of absolute differences in factor scores in each of the five personality dimensions (the more similar each social bond pair, the smaller the difference values), on the response variable social bonds, i.e. DSI scores. Due to the expected effect of absolute dominance rank differences on DSI, we included it as fixed effect. Since group composition changed between years, the same groups in the two consecutive years were handled separately, so we included a combined variable ‘group.year’ as fixed effect with 7 levels. As random effects we included ‘identity of dyad’ and ‘dominance rank difference’, calculated per half year period, controlling for the fact that they are dependent measures. Finally, random slopes were modelled for a dyads and dominance rank difference variation in DSI along ‘group.year’. We did not predict interaction effects in the model. The DSI scores were log transformed and all variables, except for ‘group.year’, were z-transformed (to a mean of zero and a standard deviation of one). The model was fitted in R (R Core Team 2017) using the function ‘lmer’ of the R-package ‘lme4’ (version 1.1-15; Bates et al., 2014).

Our visual inspection of a qq-plot, and the residuals plotted against fitted values, did not reveal obvious deviations from the model assumptions of normally distributed and homogeneous residuals.

The function ‘vif’ of the R-package ‘car’ (Fox & Weisberg, 2011; applied to a standard linear model excluding the random effects) indicated collinearity to be no issue (largest VIF=1.13; (Fidell & Tabachnick, 2003; Field, 2000; Quinn & Keough, 2002; Zuur, Ieno, & Elphick, 2010). We tested the full against the null model, comprising only ‘group.year’ as fixed effect and the random effects as described above. We fitted both models using Maximum Likelihood (rather than Restricted Maximum Likelihood; Bolker et al., 2009) and conducted a likelihood ratio test (R-function ‘anova’ with argument test set to “Chisq”; Dobson, 2010; Forstmeier & Schielzeth, 2011). To extract p-values for the individual effects, we used the R-function ‘drop1’ (with argument test set to “Chisq”; Barr, Levy, Scheepers, & Tily, 2013), based on likelihood ratio tests comparing the full to respective reduced models. Confidence intervals (lower: 2.5%, upper: 97.5%) for the estimates were computed with the function ‘confint.merMod’ of the R-package ‘lme4’ (version 1.1-15; Bates et al., 2014). The sample size for this model was a total of 140 observations made on 101 dyads and 40 absolute dominance rank differences.

We tested for potential circularity problems arising from using the same behavioural variables (body contact, grooming and friendly approach) to assess personality structure, as well as affiliative relationship strength (DSI). In case of a circularity issue, on the one hand we would expect a strong positive Pearson correlation between the two measures across individuals. We correlated the individual personality scores with the sum DSI of the top two social bond partners for each individual. On the other hand, across dyads we would expect a strong positive correlation of DSI and the mean of both partners’ personality scores on a social dimension. Pearson correlations with individual and dyadic Connectedness and Sociability scores were performed for each half year period.

To assess whether males adjusted their personality after migrating into a new group with new partners, we correlated each of the variables loading high on Connectedness (as quantified from the two-year data collection period; Table 2) across the six migrating males from one year to the next. We used Pearson correlation and variables were aggregated for April 2014– March 2015 and April 2015–March 2016.

### Ethical statement

Our animal research was completely non-invasive and approved by the Department of National Parks, Wildlife and Plant Conservation (DNP), Thailand (permit 0002/2424). This work followed the ASAB guidelines for the treatment of animals in behavioural research and teaching, and adhered to standards as defined by the European Union Council Directive 2010/63/EU on the protection of animals used for scientific purposes.

## RESULTS

The full model describing variation in dyadic relationship strength was significantly different from the null model (likelihood ratio test: χ^2^ = 14.69, df = 6, P < 0.05). The Connectedness score (likelihood ratio test: χ^2^ = 5.14, df = 1, P = 0.023) and the dominance rank difference (likelihood ratio test: χ^2^ = 4.11, df = 1, P = 0.043) had significant effects on social bonds (Table 2; Fig. 1 and 2). In accordance with previous findings, that closely bonded individuals pull each other to similar ranks (Schülke et al., 2010), we found that bond strength was associated with similarity in dominance rank. The smaller the Connectedness score of a dyad, i.e. the more similar two partners are in that personality dimension, the higher the DSI score, i.e. the stronger the social bond. Since all variables entered into the model were z-standardized, the results can be interpreted as follows: if the absolute difference in the Connectedness score of a dyad increases one standard deviation then social bond strength will decrease by about 0.18 standard deviations, with all other control variables held on average. In other words, if the Connectedness score of a dyad increases one unit then social bond strength will decrease about 0.09 units.

The graph shows that with high difference scores in Connectedness, the DSI of a dyad is far below the meaningful threshold of one, which marks strong social relationships (i.e. social bonds). The raw data are quite scattered probably due to the relatively small sample size and relatively short period to measure the social bond strength. We pooled data from four different study groups and two time periods. These were rather stable periods within an unstable observation period with alpha rank changes and migration events, which are influencing the social bonds of all group members. However, the narrow confidence intervals of the model prediction are indicative of reliable results. The personality effects are rather small like in the other primate studies (effect range: |0.043–2.02|; Capitanio et al., 2015; Massen & Koski, 2014; Morton et al., 2015; Weinstein & Capitanio, 2012) as well as in humans (Feiler & Kleinbaum, 2015; Jensen-Campbell et al., 2002; Roberts, Kuncel, Shiner, Caspi, & Goldberg, 2007).

**Table 2.** Effects of personality similarity on the strength of dyadic social bonds. Bond strength is the log standardized dyadic composite sociality score (DSI), and similarity in each of five personality dimensions was modelled as the absolute difference in personality scores between partners and dominance similarity as absolute dominance rank difference. All variables z-transformed. Significant results marked in bold.

| Variable | Estimate | SE | CI<sub>lower</sub> | CI<sub>upper</sub> | $\chi^2$ | Df | P |
| --- | --- | --- | --- | --- | --- | --- | --- |
| (Intercept) | -0.01 | 0.16 | -0.33 | 0.31 | (1) | (1) | (1) |
| Aggressiveness<sub>BC</sub> score<sup>(2)</sup> | -0.09 | 0.08 | -0.24 | 0.07 | 1.32 | 1 | 0.251 |
| Confidence<sub>TR</sub> score<sup>(3)</sup> | -0.02 | 0.08 | -0.19 | 0.16 | 0.03 | 1 | 0.853 |
| Connectedness<sub>BC</sub> score<sup>(4)</sup> | **-0.18** | **0.08** | **-0.33** | **-0.02** | **5.14** | **1** | **0.023** |
| Sociability<sub>BC</sub> score<sup>(5)</sup> | -0.03 | 0.08 | -0.19 | 0.13 | 0.14 | 1 | 0.706 |
| Vigilance<sub>BC</sub> score<sup>(6)</sup> | -0.10 | 0.08 | -0.26 | 0.05 | 1.77 | 1 | 0.183 |
| Dominance rank difference<sup>(7)</sup> | **-0.19** | **0.09** | **-0.37** | **-0.01** | **4.11** | **1** | **0.043** |
| Group.year | (8) | (8) | (8) | (8) | 3.81 | 6 | 0.702 |

<sup>(1)</sup>not shown, because having a limited interpretation.
<sup>(2)</sup>z-transformed, original values with mean ± SD: $1.18 \pm 0.79$
<sup>(3)</sup>z-transformed, original values with mean ± SD: $1.24 \pm 0.85$
<sup>(4)</sup>z-transformed, original values with mean ± SD: $1.01 \pm 0.70$
<sup>(5)</sup>z-transformed, original values with mean ± SD: $1.10 \pm 1.10$
<sup>(6)</sup>z-transformed, original values with mean ± SD: $1.12 \pm 0.92$
<sup>(7)</sup>z-transformed, original values with mean ± SD: $2.98 \pm 1.88$
<sup>(8)</sup>7 levels of group.year reveal no effect and are not shown.

**Figure 1:**
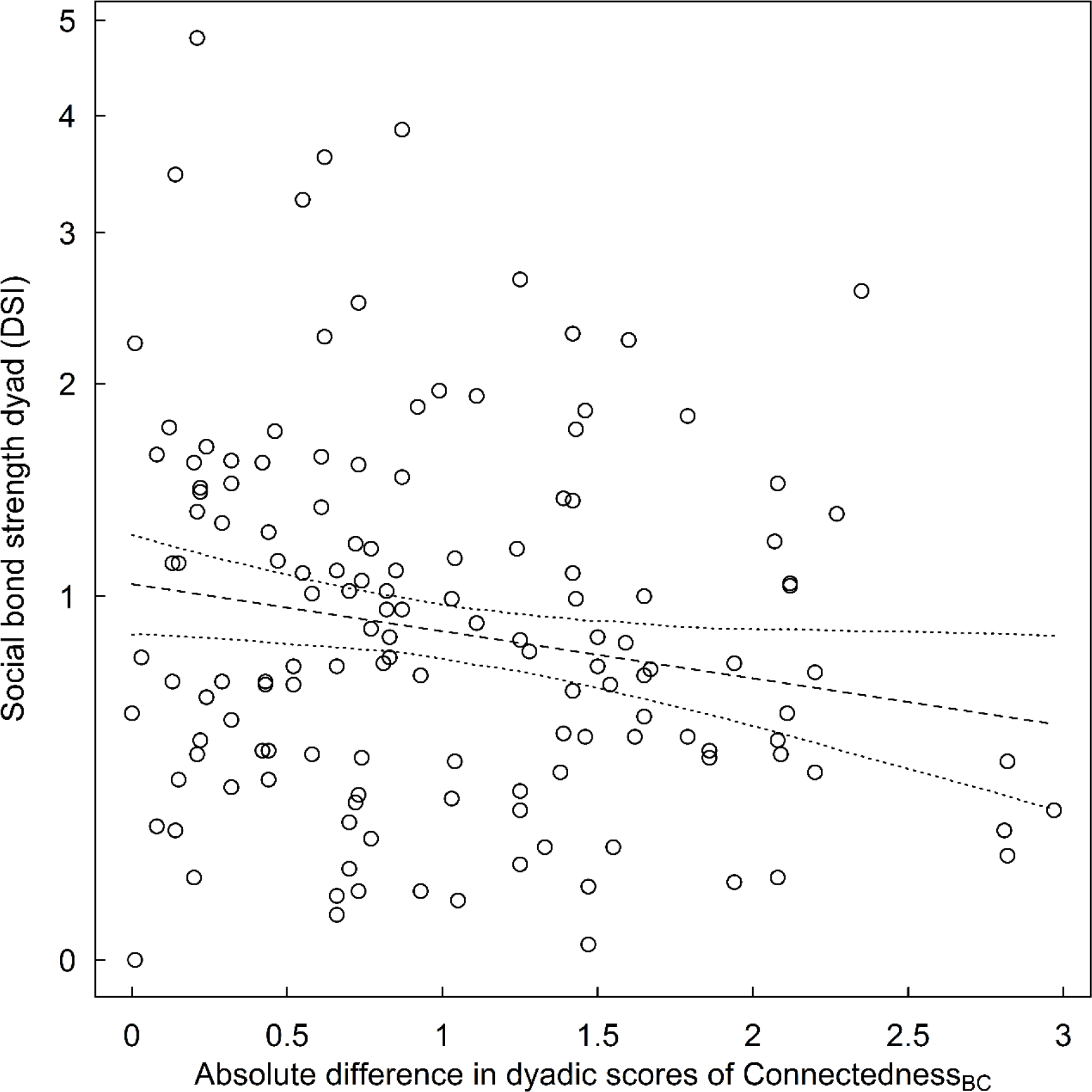
Effect of personality similarity on bond strength. Linear mixed model plot with the significant effect of absolute difference in dyadic scores of Connectedness on log standardized social bond strength (DSI). The dashed line is the model prediction and dotted lines represent its bootstrapped 95% confidence intervals. Total N with 101 dyads and 40 dominance rank differences. All variables z-transformed.

**Figure 2:**
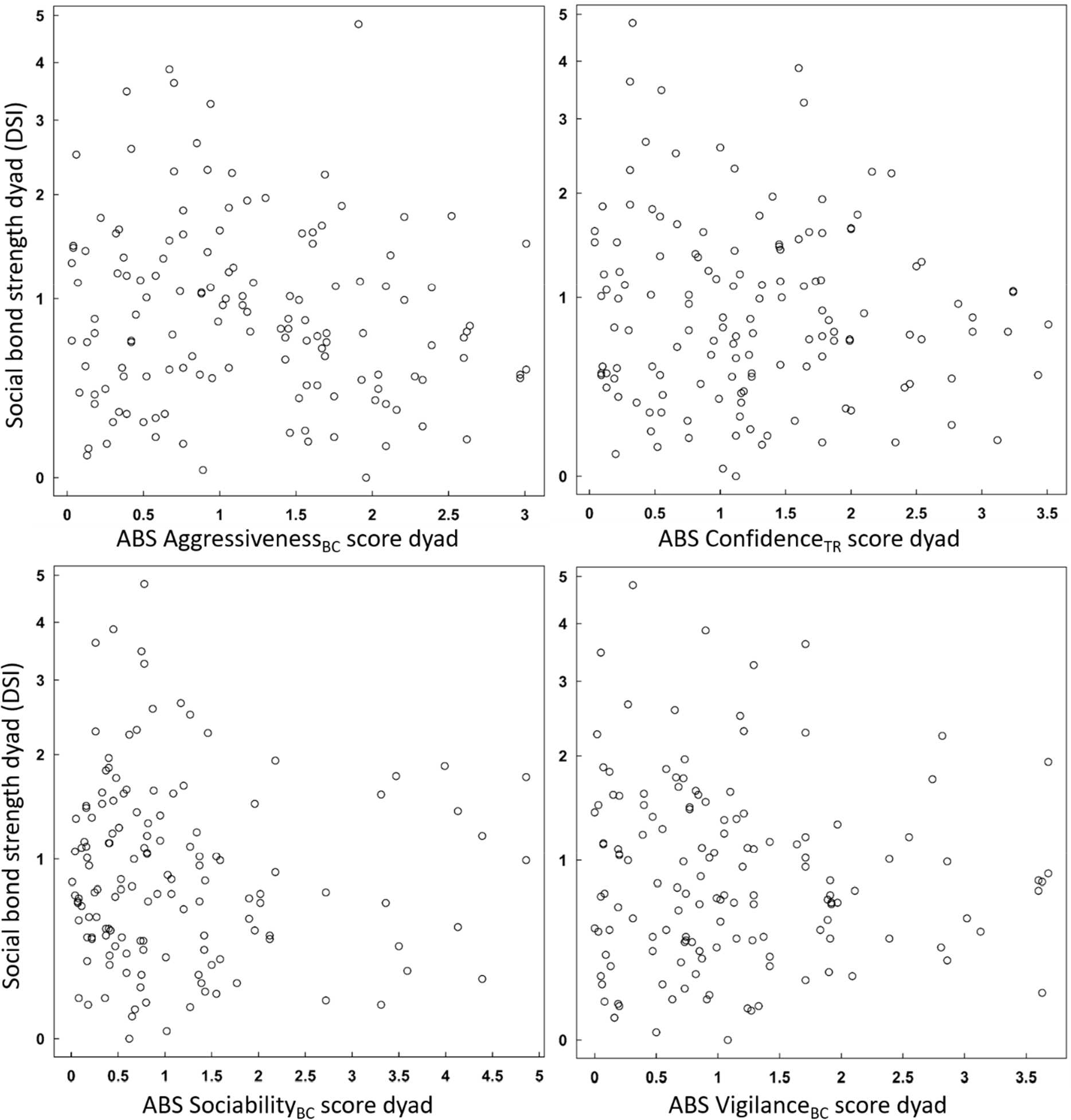
Other personality traits (absolute difference (ABS) in dyadic scores) with no effect on social bond strength (log standardized DSI). Total N with 101 dyads and 40 dominance rank differences. All variables z-transformed.

The strength of affiliative relationships was explicitly related to the similarity in personality between partners and did not result from dyads or individuals scoring high or low on social personality dimensions. DSI did not correlate with mean Connectedness of a dyad (Oct2014–Mar2015: *r*_dyadic_= 0.139, *p*=0.227, n=77; Oct2015–Mar2016: *r*_dyadic_=0.054, *p*=0.675, n=63) and mean Sociability scores per dyad (Oct2014–Mar2015: *r*_dyadic_=−0.135, *p*=0.242, n=77; Oct2015–Mar2016: *r*_dyadic_ = 0.246, *p* = 0.052, n = 63; Fig. A1). Similarly, the strength of the strongest bonds this individual formed (i.e. sum of top two DSI values) did not correlate with individual Connectedness (Oct2014–Mar2015: *r*_individual_ = 0.076, *p* = 0.722, n = 24; Oct2015–Mar2016: *r*_individual_ = −0.004, *p* = 0.985, n = 21) and Sociability scores (Oct2014–Mar2015: *r*_individual_ = −0.155, *p* = 0.471, n = 24; Oct2015–Mar2016: *r*_individual_ = 0.168, *p* = 0.468, n = 21; Fig. A2).

### Friendship formation

For our small subset of six migrating individuals, the variables loading on the Connectedness dimension were positively correlated from before to after the migration for variables active, alone, neighbour diversity and tolerance (mean *r* = 0.817; *p* = 0.02–0.1; Table 3), with the exception of friendly approach (*r* = 0.041; *p* = 0.94; Table 3).

**Table 3.** Stability in variables loading on the Connectedness personality domain in six males that changed groups.

| Behavioural variable | Pearson's $r$ | $p$-value |
| --- | --- | --- |
| active | 0.879 | 0.02 |
| alone | 0.724 | 0.10 |
| friendly approach | 0.041 | 0.94 |
| neighbour diversity | 0.860 | 0.03 |
| tolerance | 0.805 | 0.05 |

## DISCUSSION

Consistent with the idea that partner choice in social bond formation is guided by personality homophily, male Assamese macaques chose bond partners with similar levels of Connectedness. Similarity in Connectedness most likely predicted social bond formation and not the other way around, because males did not change their personality after migrating to a new group. In the following we will compare these results with personality homophily in humans, discuss its adaptive value, evidence from animal mating pairs and other types of animal social bonds, and why partners are similar in social personality traits and not in other dimensions. We discuss the role of tolerance in bonding and cooperation and their neural basis and consider alternative theories for the social effects of partners’ personality.

Our result that individuals more similar in Connectedness form stronger social bonds supports the hypothesis of a shared evolutionary origin of personality homophily as partner choice strategy in human and non-human primates (Bahns et al., 2016; Massen & Koski, 2014). In humans the personality dimensions most closely matched in friends are extraversion and agreeableness (e.g., Blaz, 1983; Caspi et al., 2005; Dishion, Patterson, Stoolmiller, & Skinner, 1991; Ilmarinen, Vainikainen, Verkasalo, & Lönnqvist, 2017; Maaß, Lämmle, Bensch, & Ziegler, 2016; Markey & Kurtz, 2006; van Zalk & Denissen, 2015; Youyou et al., 2017), which partly resembles our findings. Aspects of the Connectedness trait, like proximity, social tolerance, and friendly approach, roughly correspond to the sociable or affiliative facets of extraversion associated with enjoyment of social interactions (Denissen & Penke, 2008). Our sociability domain (i.e. high rates of friendly behaviour and more time in body contact and grooming, as well as more frequent initiation of affiliations) has more overlap with the agreeableness dimension in humans, where individuals scoring higher in agreeableness are more interested in maintaining positive social relationships (Denissen & Penke, 2008). Unlike in humans, homophily regarding this second social personality dimension did not predict social bonds.

The main selective advantage of personality similarity in friendships as well as animal social bonds may result from a more reliable and thus more successful cooperation among individuals with similar (cooperative) behavioural tendencies via facilitated coordination, communication and reciprocity, as well as reduced uncertainty and conflict (Asakawa-Haas, Schiestl, Bugnyar, & Massen, 2016; Bahns et al., 2016; Chiang & Takahashi, 2011; Curry & Dunbar, 2013; Fu et al., 2012; Gabriel & Black, 2012; Koski & Burkart, 2015; Massen & Koski, 2014; Riolo et al., 2001; Schuett et al., 2011). Humans cultivate cooperative relationships sustained by emotional closeness and reciprocity of support (Dunbar, 2018; Hruschka & Henrich, 2006; Rand & Nowak, 2013; Wrzus & Neyer, 2016)(Dunbar, 2018; Hruschka & Henrich 2006; Rand & Nowack, 2013; Wrzus & Neyer, 2016), whereby people preferentially form ties with others who share similar cooperative behavioural tendencies (Apicella et al., 2012). Extraversion and agreeableness are linked to motivation for cooperative activities as well as cooperative skills. For instance, people scoring high in these dimensions have greater enthusiasm toward cooperation and are more trusting of others (Adali & Golbeck, 2012; Ashton, Paunonen, Helmes, & Jackson, 1998; Hirsh & Peterson, 2009; Lu & Argyle, 1991; Ross, Rausch, & Canada, 2003; but see also: Koole, Jager, van den Berg, Vlek, & Hofstee, 2001).

Animal mating pairs of partners with similar level in exploration tendency (rodents: Rangassamy et al., 2015; Steller‘s jays: Gabriel & Black, 2012; great tits: Dingemanse et al., 2004; zebra finches: Schuett et al., 2011) and boldness (guppies: Ariyomo & Watt, 2013) express higher reproductive success, and successful cooperative-breeding common marmosets show group-level similarity in both traits (Koski & Burkart, 2015). The role of similarity in social personality traits remains underexplored. Exploration may be more directly linked to helping behaviour, as demonstrated in a cooperative-breeding cichlid (Bergmüller & Taborsky, 2007) and choices for breeding partners may differ in choices for other partnerships where social personality traits may be more relevant (Koski, 2014).

Across group members, chimpanzees and Capuchin monkeys show proximity driven, i.e. social tolerance related, personality homophily in social relationships (Massen & Koski, 2014; Morton et al., 2015). Further, in a trait rating study with juvenile rhesus macaques, an equitability dimension (e.g., calmer, more easygoing, less active), which also includes aspects of social tolerance, correlated with relationship stability (Weinstein & Capitanio, 2012). However, in a social network study with wild Barbary macaques, it was not similarity in social tolerance but excitability (contains elements related to low impulse control: excitable, impulsive, erratic and disorganized) that was correlated with spatial association (Tkaczynski, 2017), albeit this effect was not seen in grooming networks.

More generally, social tolerance (i.e. tolerating the proximity of others), as well as social grooming behaviour, are considered as prerequisites for animal social bonds, and, like friendships, they are further assumed to require mutuality and positive interactions (Asakawa-Haas et al., 2016; Brosnan et al., 2015; Massen, Sterck, & De Vos, 2010; van Zalk & Denissen, 2015; Watts, 2002). Considering homophily in Connectedness as partner choice mechanisms in Assamese macaques, similar needs of proximity and similar level of social tolerance (scoring either high or low in Connectedness), may be associated with increased trust in reciprocal relations with bond partners, to maintain bonds and facilitate cooperation (Campennì & Schino, 2014; Laakasuo, Rotkirch, Berg, & Jokela, 2016; Massen & Koski, 2014). Cooperative success and bond maintenance are intertwined regarding social bonds as alliances that generate adaptive benefits via support in critical situations (DeScioli & Kurzban, 2009; Massen & Koski, 2014; Schülke et al., 2010; Seyfarth & Cheney, 2012). Mutual coalitionary support helps bond partners to attain and maintain high social status, which is linked to reproductive success in male Assamese macaques (Schülke et al., 2010; Sukmak, Wajjwalku, Ostner, & Schülke, 2014). In Barbary macaques, it was demonstrated experimentally that strong social bonds positively influenced the maintenance of cooperation over a long period (Molesti & Majolo, 2016).

Social tolerance (or other traits in other species) may be correlated with cooperativeness, given that correlations between different behaviours are assumed to occur among different functional contexts (behavioural syndromes: Bergmüller, Schürch, & Hamilton, 2010; Sih, Bell, & Johnson, 2004; see also cooperative syndromes in cooperative breeding meerkats: Clutton-Brock, Russell, & Sharpe, 2003; English, Nakagawa, & Clutton-Brock, 2010 and cichlids: Schürch & Heg, 2010). Social tolerance could as well be functionally related to variation in other cognitive abilities or styles to negotiate the social landscape, which in turn affect cooperation (Fiske & Haslam, 1996; Moreira et al., 2013; Seyfarth & Cheney, 2015; Sih & Del Giudice, 2012). Differences in social awareness or sensitivity, comprising the ability to monitor the cooperative tendencies of others, may favour the evolution of consistent individual differences in cooperation (Korman, Voiklis, & Malle, 2015; McNamara, Stephens, Dall, & Houston, 2009; Seyfarth & Cheney, 2015; cognitive syndromes: Sih & Del Giudice, 2012). It was recently demonstrated that chimpanzees high in Extraversion (corresponding to Assamese’ Connectedness) and assumingly more sensitive to inter-individual interactions, have been more sensitive to inequity in outcomes between themselves and a social partner in an experimental condition (Brosnan et al., 2015). In sum, homophily in social tolerance in Assamese macaques may either be related to similar cooperative tendencies or similar social sensitivity in bonded partners leading to enhanced cooperative success, probably because of increased trust in compatible partner.

Friends show similar neural responses to the same stimuli and thus react to the world around them in a similar way, presumably due to similar dispositions, pre-existing knowledge, opinions, interests, and values (Parkinson et al., 2018). Such similar neural responses are proposed to enhance social interactions and friendship formation via positive affective processes, increased predictability and facilitated communication (Berger & Calabrese, 1975; Neyer et al., 1999; Selfhout et al., 2010; van Zalk & Denissen, 2015). The same line of argument may apply to animal social bonds. Similarity in personality, or possibly social tolerance traits in particular, may trigger basic neural and physiological mechanisms (underlying social interactions in humans and other animals: e.g., Brent 2014; Chang et al., 2013; Dunbar, 2010), in the bond partner in a similar way, which in turn may facilitate attitudinal or emotionally based partner choice (Fruteau, Voelkl, Van Damme, & Noë, 2009; Fu et al., 2012; Parkinson et al., 2018; Schino & Aureli, 2009). Koski & Burkhart (2015) propose that similar affective states may facilitate behavioural synchrony, contingency and reciprocity in a cognitively inexpensive way (Brosnan & de Waal, 2002; Fessler & Holbrook, 2014). Not alone that long-term relationships may be reliably maintained via emotionally based reciprocity (Schino & Aureli, 2016), positive affect and common psychological mechanisms may allow for quick assessment in bond formation as well, since it is known in humans that similar people relate with each other quite rapidly and without concise choice (Ambady, Bernieri, & Richeson, 2000; Bahns et al., 2016; Sunnafrank & Ramirez, 2004).

Alternative theories in human personality research claim that ‘opposites attract’. Interpersonal theory (Carson, 1969), proposes that dominance invites submission and vice versa, while partners mutually reinforce each other’s dispositional tendencies. Self-expansion theory (Aron & Aron, 1996) suggests that people accommodate to each other’s distinctiveness to expand their selves. Empirical studies often found mixed evidence. For instance, friends were either very similar or very different regarding extraversion-introversion (Nelson et al., 2011). Pairings of rhesus macaques in a laboratory setting were successful for females similar in Emotionality, but only for those males with both dyad members scoring low (but not moderate or high) on Gentle and Nervous temperament (Capitanio et al., 2015). Yet, researchers mostly agree that homophily plays an important role in long-term relationships. When people form relationships with dissimilar individuals these are rather short-lived task-oriented ties, like professional collaborations (Currarini, Jackson, & Pin, 2009; Fu et al., 2012; McPherson et al., 2001; Moody, 2004; Parkinson et al., 2018; Rivera et al., 2010).

Another alternative theoretical account for the observed correlations between personality and social relationships invokes social influence and predicts that friends may become more similar over time, and individuals may potentially converge their attitudes to one another to be more liked (normative) or to be more right (informational) (Cullum & Harton, 2007; Davis & Rusbult, 2001). Likewise, there is evidence for post pairing adjustment (associated with improved reproductive success) with reactive partners becoming more proactive in monogamous fish (Laubu, Dechaume-Moncharmont, Motreuil, & Schweitzer, 2016). Consistent with our finding that personality similarity most likely predicts social bond formation in Assamese macaques, human studies demonstrated that similarity matters early in acquaintanceship, and established attitudes, values and personality seem generally less amenable to influence (Bahns et al., 2016; Costa & McCrae, 1992; Papadopoulou, 2016). Still, not many studies considered social influence, and further research is needed especially in the realm of animal social bonds.

In fact, human psychology research even goes beyond the statement of selectivity in friendships, and proposes that people engage in niche construction when they seek out social environments, such as friendships (e.g., Kandel, 1978; Bahns et al., 2016; Papadopoulou, 2016). In short, Niche Construction Theory (NCT; Odling-Smee, Laland, & Feldman, 2003), refers to evolutionary processes as constant and cyclical transactions between the organisms, their socio-physical environment and their genetic heritage, whereby organisms modify their own (and/or each other’s) environments through the metabolic, physiological and behavioural activities, as well as through their choices (Flynn, Laland, Kendal, & Kendal, 2013; Laland, Odling-Smee, & Endler, 2017; Odling-Smee et al., 2013). Recent studies investigated friendship dyads in adults and children in a real-life setting, and newly formed relationships were tracked over some period (Bahns et al., 2016; Papadopoulou, 2016). These studies support previous findings and state that humans actively choose similar minded (e.g., on personality or attitudes) friends to construct stable, satisfying social niches, that are compatible with their dispositions, and further promote cooperation and well-being (Bahns et al., 2016; Caspi & Herbener, 1990; Hampson, 2011; Papadopoulou, 2016; Scarr & McCartney, 1983).

In sum, our results support the idea of a fundamental biological basis of homophily as partner choice strategy in human and non-human animals (Apicella et al., 2012; Bahns et al. 2016; Fu et al., 2012; Massen & Koski, 2014). Specifically, homophily in social tolerance traits may play an important role considering the potential relatedness of human personality traits extraversion and agreeableness with the Connectedness domain in Assamese macaques plus the evidence from other primate studies relating personality and social bonds (Massen & Koski, 2014; Morton et al., 2015; Weinstein & Capitanio, 2012). Further, social tolerance is key in social bonds and cooperative success (e.g., raven: Asakawa-Haas et al., 2016; Massen et al., 2015; hyena: Drea & Carter, 2009; primates: Hare, Melis, Woods, Hastings, & Wrangham, 2007; Werdenich & Huber, 2002; theoretical model: Chen, Fu, & Wang, 2009). To gauge the generality of these findings, additional primate and particularly other animal studies are needed to elucidate the importance of similarity in social tolerance in the process of social bond formation.

## ACKNOWLEDGMENTS

We thank the National Research Council of Thailand (NRCT) and the Department of National Parks, Wildlife and Plant Conservation (DNP) for permission to conduct this study and for all the support granted (permit 0002/2424). We are grateful to I. Chatchawarn, J. Prabnasuk, K. Nitaya T. Wongsnak, M. Pongjantarasatien and K. Kreetiyutanont, M. Kumsuk, W. Saenphala (PKWS) for their cooperation over the years and permission to carry out this study. We thank A. Koenig and C. Borries, who developed the field site. Our special thanks goes to S. Jumrudwong, W. Nueorngshiyos, N. Juntuch, J. Wanart, R. Intalo, T. Kilawit, N. Pongangan, B. Klaewklar, N. Bualeng, D. Gutleb, P. Saisawatdikul, K. Srithorn, M. Swagemakers and T. Wisate for their excellent help in the field, and to P. Saisawatdikul and C. Haunhorst for their support. We thank Roger Mundry for the permission to use statistical packages developed by himself. This research was funded by the Deutsche Forschungsgemeinschaft (DFG, German Research Foundation) – Project number 254142454 / GRK 2070.

## APPENDIX

**Table A1.** Group composition with age-sex classes at onset of study.

| Group | Adult<br>males | Subadult<br>males | Juvenile<br>males | Infant<br>males | Adult<br>females | Juvenile<br>females | Infant<br>females | Total |
| --- | --- | --- | --- | --- | --- | --- | --- | --- |
| ASM | 8 | 6 | 10 | 3 | 10 | 12 | 3 | 52 |
| AOM | 10 | 3 | 8 | 1 | 14 | 10 | 5 | 51 |
| ASS | 4 | 2 | 2 | 4 | 9 | 7 | 1 | 28 |
| AOS | 5 | 0 | 2 | 2 | 6 | 5 | 1 | 21 |
| All | 27* | 11 | 22 | 10 | 39 | 34 | 10 | 153 |

\*24 adult males were included in the analysis because 3 individuals emigrated.

**Figure A1:**
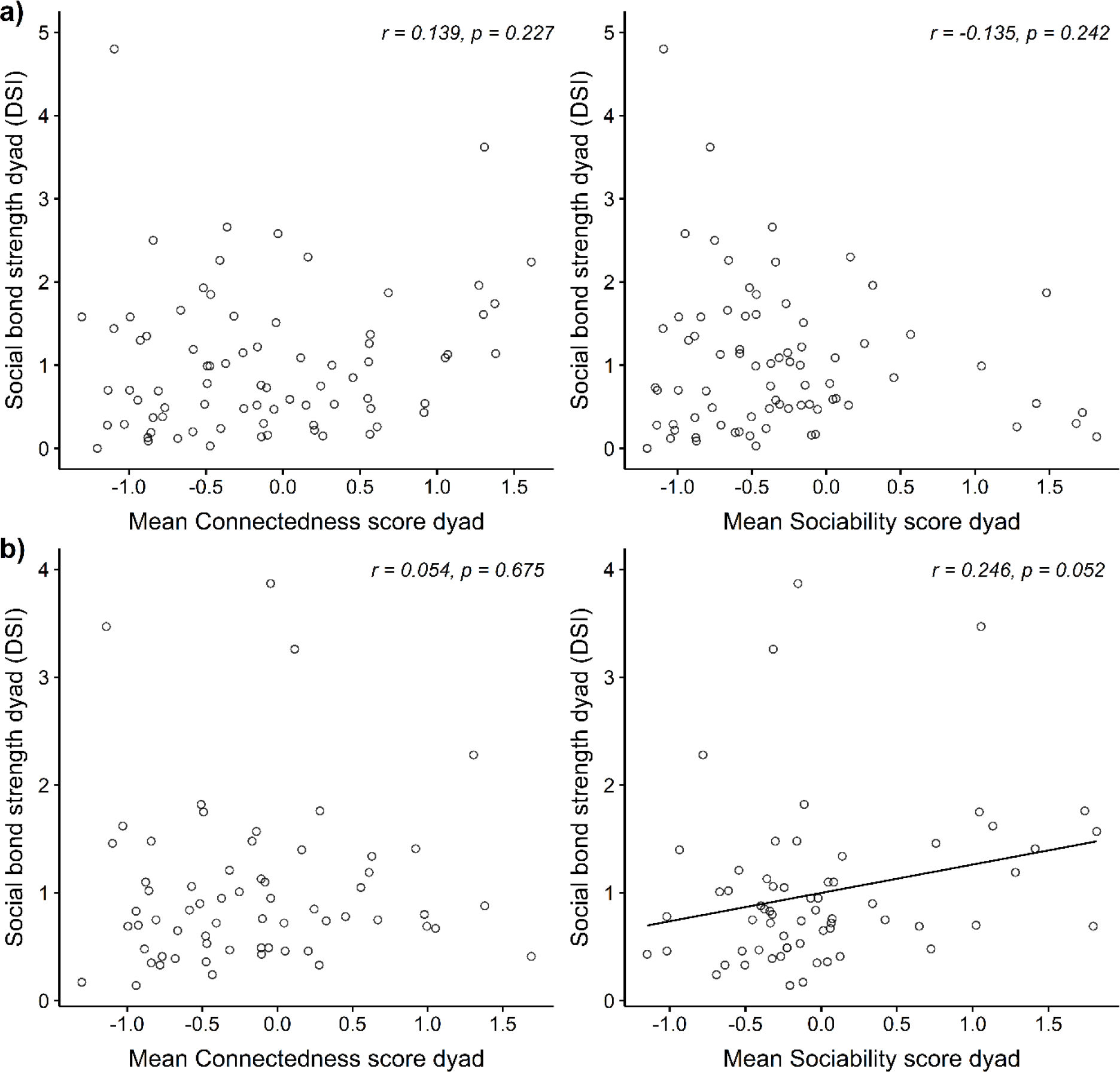
Pearson correlation of mean personality scores per dyad with DSI scores for every half year period. a) Oct2014–Mar2015 with n=77. b) Oct2015–Mar2016 with n=63.

**Figure A2:**
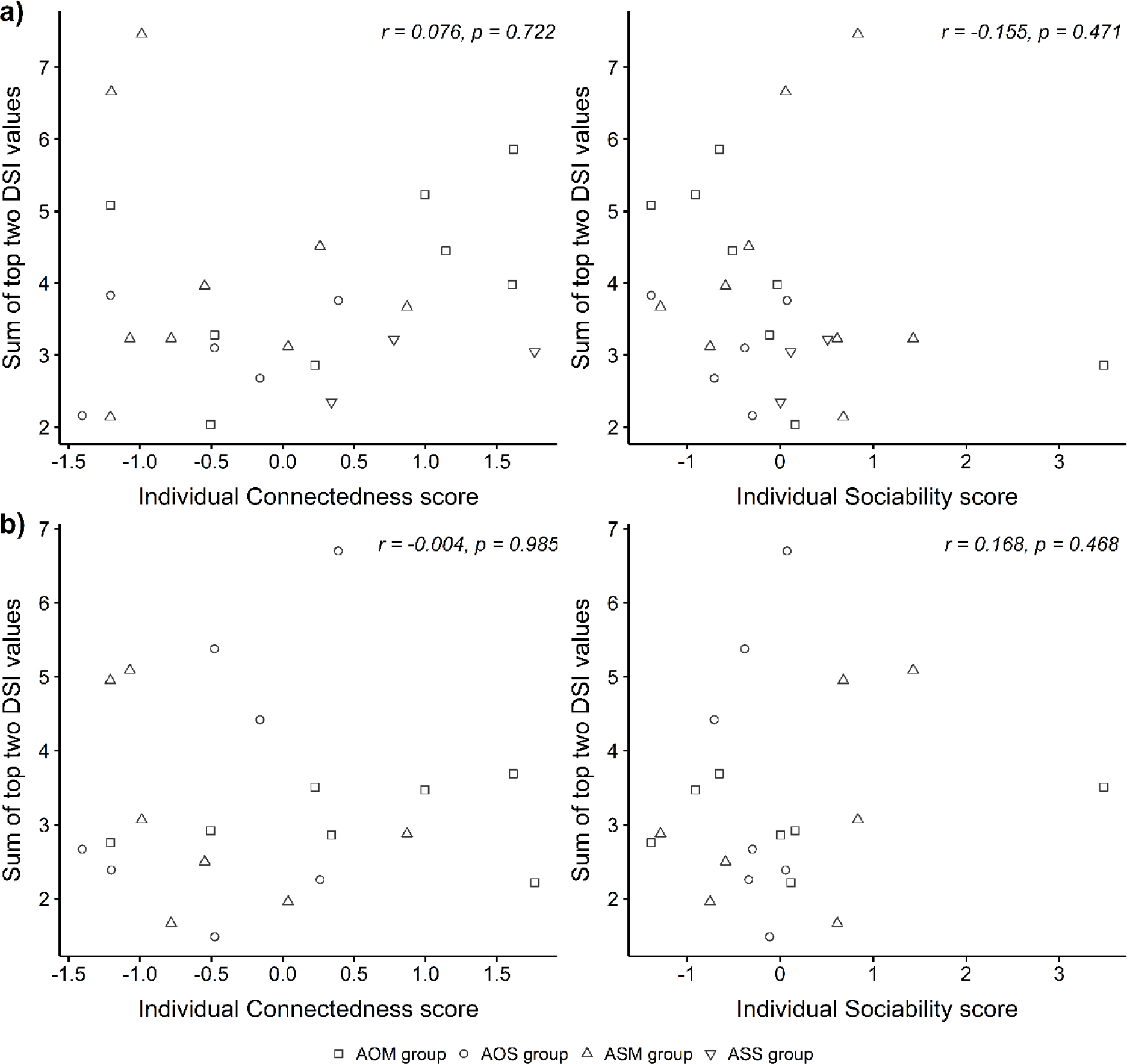
Pearson correlation of individual personality scores with sum of top two DSI values for every half year period. a) Oct2014–Mar2015 with n=24. b) Oct2015–Mar2016 with n=21.

